# Effects of orally administered crofelemer on the incidence and severity of neratinib-induced diarrhea in female dogs

**DOI:** 10.1101/2023.02.23.529666

**Authors:** Michael K. Guy, Andre Teixeira, Allison Shirer, James Bolognese, Pravin Chaturvedi

## Abstract

Management guidelines for cancer therapy-related diarrhea (CTD) should be revised because newer targeted therapies have increased CTD burden, with high incidence and/or severity of diarrhea for some agents that inhibit epidermal growth factor receptor and receptor tyrosine kinases. Neratinib, a pan-HER tyrosine kinase inhibitor, approved for breast cancer treatment, causes severe diarrhea in >95% of patients. Crofelemer, a novel intestinal chloride ion channel modulator, is an approved antidiarrheal for patients with HIV receiving antiretroviral therapy. The objective of this study was to evaluate the effectiveness of crofelemer prophylaxis in reducing the incidence and severity of neratinib-induced diarrhea without loperamide in dogs. Female dogs received neratinib orally daily concomitantly with either matching placebo tablets (CTR) or crofelemer 125 mg delayed-release tablet two or four times/day (BID or QID) for 28 consecutive days. At the end of treatment, 37.5%, 75%, and 87.5% of the CTR, BID, and QID dogs were ‘responders’ defined as ≤7 loose/watery stools/week for at least 2 of 4 weeks (p<0.05). The average number of watery stools per week was 9, 6, and 6 in the CTR, BID, and QID groups, respectively (p<0.05). The average number of weeks with no loose/watery stools was 1.3, 2.1, and 2.3 for the CTR, BID, and QID groups, respectively (p<0.05). The weekly mean fecal scores and stool consistency were 5.1, 3.9, and 4.1 for the CTR, BID, and QID groups (p<0.05). In this 28-day preclinical study, crofelemer prophylaxis without loperamide reduced the incidence and severity of neratinib-associated diarrhea in female dogs by 30%.

**Ethical Compliance:** All procedures performed in studies involving canine participants were in accordance with the ethical standards of the institutional and/or national research committee and applicable Institutional Animal Care and Use Committee (IACUC).

## Introduction

Cancer therapy-related diarrhea (CTD) is a debilitating side effect in patients receiving targeted therapy, with or without chemotherapy. CTD leads to dehydration, electrolyte and fluid imbalances, renal insufficiency, malnutrition, fatigue, and other issues. CTD severely compromises cancer treatment because it requires dosing holidays and/or dose reductions of cancer drugs (**1**), resulting in ineffective cancer treatments or worse yet, resistance, and treatment failures. Furthermore, healthcare resource utilization costs are significant (**2, 3**) for cancer patients with diarrhea. CTD occurs in 50-80% of patients depending on the targeted therapy and/or chemotherapy regimens, and the advent of newer targeted therapies has increased the burden of gastrointestinal toxicity of these drugs because of their mechanism of action, specifically for those that target epidermal growth factor receptor (EGFR), tyrosine kinases, and human epidermal growth factor receptor 2 (HER2) (**4**). Tyrosine kinase inhibitors (TKIs), such as neratinib, result in a very high incidence of diarrhea (>95%). The management of CTD to reduce and/or prevent the severity and duration of diarrhea is critically needed.

Crofelemer is a novel antisecretory, antidiarrheal, oral botanical drug purified from the crude plant latex of *Croton lechleri*. Crofelemer elicits its effects on apical membrane transport and signaling processes involved in intestinal chloride ion transport. Crofelemer reduces chloride ion secretion by the Cystic Fibrosis Transmembrane conductance regulator (CFTR) channel through use-dependent inhibitory modulation of CFTR, resulting in an IC_50_ of 6-7 μM (**5**). Crofelemer’s action resisted washout, with inhibition lasting several hours after washout. Crofelemer was also found to strongly inhibit the intestinal calcium-activated chloride channel TMEM16A, also known as ANO1 and DOG (anoctamin 1 and Discover On Gastrointestinal Stromal Tumors) by a voltage-independent inhibition mechanism with a maximum inhibition of 90% and an IC_50_ of 6.5 μM (**5**). The dual inhibitory action of crofelemer on two structurally unrelated 5 intestinal Cl^−^ ion channels results in its unique and novel physiological antisecretory antidiarrheal effects in reducing fluid and electrolyte accumulation in the gastrointestinal lumen and improving stool consistency.

Crofelemers are approved by the US FDA for the symptomatic relief of noninfectious diarrhea in adult HIV/AIDS patients on antiretroviral therapy. Crofelemers have demonstrated safety and efficacy in people living with HIV/AIDS in numerous clinical studies when compared with placebo in reducing the frequency of loose/watery bowel movements as well as improvement in stool consistency (**6, 7, 8, 9**).

Neratinib is an oral irreversible HER1, HER2, and HER4 tyrosine kinase inhibitor that requires loperamide prophylaxis due to the high incidence of severe diarrhea associated with the drug (**10**). However, prophylaxis and management guidelines using antimotility drugs, such as loperamide, an opiate agonist, are empirical interventions to reduce the severity of diarrhea. Strategies for the management of CTD are needed with physiologically relevant antidiarrheal agents, which would correct stool consistency, rather than reduce gastrointestinal motility, preferably as prophylaxis, as the chronic and/or prophylactic use of antimotility drugs has been inadequate or ineffective.

The objective of this study was to evaluate the prophylactic effects of crofelemer in female dogs experiencing neratinib-induced diarrhea without loperamide prophylaxis, following concomitant neratinib and crofelemer (twice or four times daily) or placebo administration for 28 consecutive days.

## Materials and Methods

### Animals

Twenty-four female beagle dogs (*Canis familiaris*) between 8 and 9 months of age, weighing 6.4 – 8.5 kg were acquired (Marshall Farms, North Rose, NY, USA). All study participants underwent quarantine and acclimation at the study site for 21 days. During the acclimation period, the dogs were observed at least once daily for any abnormal clinical signs. All dogs had normal behavior and were randomized into the study.

### Study Design

Twenty four (24) healthy female dogs were randomized into three groups of eight dogs each receiving oral neratinib 40 mg daily, with one group of eight dogs receiving oral neratinib 40-80 mg with crofelemer placebo tablets every 6 h (Control – CTR group), a second group of eight dogs receiving oral neratinib 40-80 mg with crofelemer 125 mg delayed-release tablets every 12 h (BID group), and a third group of eight dogs receiving oral neratinib 40-80 mg with crofelemer 125 mg delayed-release tablets every 6 h (QID group). Neratinib tablets (Nerlynx 40 mg) were provided by Puma Biotechnology, Inc., and Crofelemer 125 mg delayed-release tablets (Mytesi) were provided by Napo Pharmaceuticals Inc.

### Enrollment and Dose Administration

All 24 dogs in this study had diarrhea induced by neratinib, as evidenced by the passage of liquid stool (fecal score of 6 or 7). The daily dose of neratinib was equivalent to the human dose of 5-10 mg/kg/day. Because of emesis in the first few days of dosing, the dogs were initially administered one 40 mg tablet of neratinib orally once daily for the first 5 days (5 mg/kg/day), and then the daily oral dose of neratinib was increased to two tablets (80 mg) once daily (10 mg/kg/day) for the remainder of the 28-day study period. Neratinib was titrated daily to maintain diarrhea without severe signs of inappetence, lethargy, dehydration, vomiting, and bloody diarrhea. The crofelemer or placebo-controlled treatments for all dogs started on the same morning as the first dose of neratinib, and the study treatments continued for 28 consecutive days.

### Data Collection

Body weight, oral fluid administration (based on the clinical evaluation of hydration), and qualitative food consumption were recorded daily. Fecal scores were collected twice daily for 28 consecutive days in all dogs across the three treatment groups. The Purina Fecal Scoring (PFS) system, which uses a seven-point scale analogous to the human Bristol Stool Form Scale, was used to assess stool consistency.

### Endpoints

The effectiveness of the crofelemer was measured by the reduction in the incidence (i.e., the number of watery bowel movements) and severity (changes in stool consistency) in dogs. The incidence of loose/watery stools was assessed using the 7-point PFS scale to distinguish loose/watery stools (PFS ≥ 6) from formed stools (PFS < 6). To assess the weekly severity of diarrhea, the Purina Fecal Score was evaluated as the average fecal score collected per observation. The following endpoints were evaluated: Responders were defined as number of dogs with ≤7 loose / watery stools (fecal scores “6” or “7”) per week for at least 2 out of the 4 weeks of the treatment period and was the primary endpoint. The average number of loose/watery stools per week, fecal scores per week, and across all four weeks were calculated. Dose reduction of neratinib (as needed on a daily basis) was monitored. Furthermore, the weekly volume of fluids required to maintain hydration was evaluated. Changes in body weight per week from baseline were evaluated.

### Statistical Methods

As the primary endpoint, bowel movements were dichotomized into watery bowel movements (fecal score of 6 or 7) or normal bowel movements (fecal score < 6). Logistic regression was used to assess the treatment effects of crofelemer compared to placebo, particularly to determine the odds of being a responder and the odds ratios versus control. A responder was defined as any dog that during the 4-week study period had at least 2 weeks of less than 7 watery bowel movements per week. For logistic regression analysis, the baseline fecal score (day 0) and baseline body weight (day 0) were used as covariates. Bowel movements were scored according to the Purina fecal score (PFS) and recorded twice daily for 28 days. Statistical tests comparing each crofelemer regimen (BID or QID) versus placebo (CTR) used a 2-sided alpha value of 0.05.

For continuous endpoints, Least Square Means (LSM, adjusted for baseline fecal scores) were analyzed using Analysis of Covariance (ANCOVA) with the respective baseline value as the covariate. Normality of residuals was assessed using the Shapiro-Wilk statistic for closeness to normality. The Shapiro-Wilk statistics for the analyses were all > 0.9, indicating that no data transformation was needed, and the untransformed summary statistics were adequate. All procedures were conducted using SAS 9.3 (Cary, NC, USA).

Two other parameters are important for an appropriate interpretation of the results: oral fluid administration (for dogs in need of oral hydration) and the dose of neratinib necessary to maintain a consistent level of diarrhea. Both variables were analyzed using a procedure similar to that described above for fecal cores for stool consistency.

The power of the study was computed using standardized effect sizes (mean difference from control divided by pooled SD). For clarity, effect sizes can be judged to be large if they are close to or greater than 1.0, as the sample sizes required for adequate power in future studies are generally small. For example, for 90% power (alpha=0.05 2-sided), N=34, 27, and 23 per treatment group would be required if the TRUE underlying effect sizes were 0.8, 0.9, and 1.0, respectively.

## Results

Induction of diarrhea by neratinib and subsequent treatment with crofelemer or vehicle control were well tolerated, as there were no deleterious effects on the dog’s behavior, body weight, hydration status, food consumption, or clinical evaluation. Concomitant daily dosing of neratinib with crofelemer or placebo did not cause any mortality during the 28-day study. In this study, 3 of 8 (37.5%) placebo-treated dogs were determined to be responders, whereas 6 of 8 (75%) crofelemer BID-treated dogs and 7 of 8 (87.5%) crofelemer QID-treated dogs were deemed responders (See **Figures 1 and 2**).

**Figure 1.**
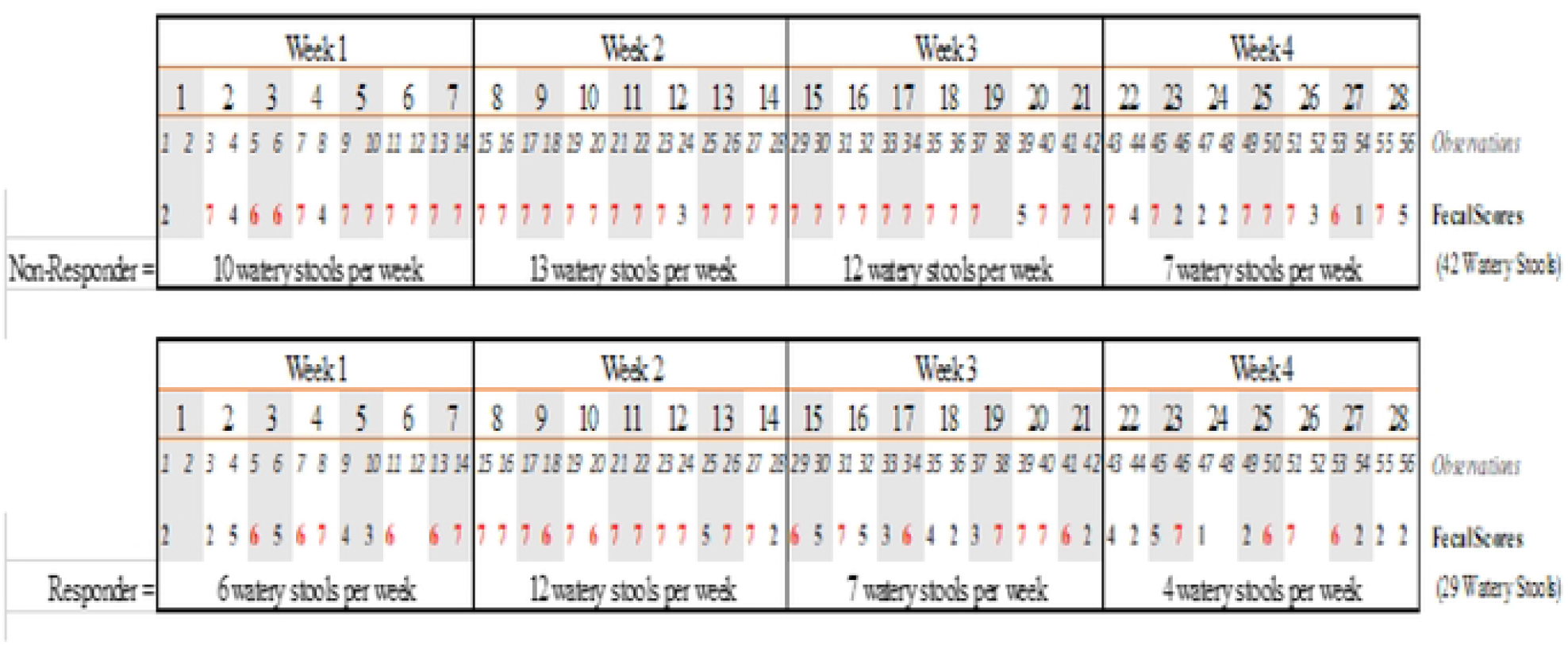
Examples of Non-Responder and Responder Dogs

**Figure 2.**
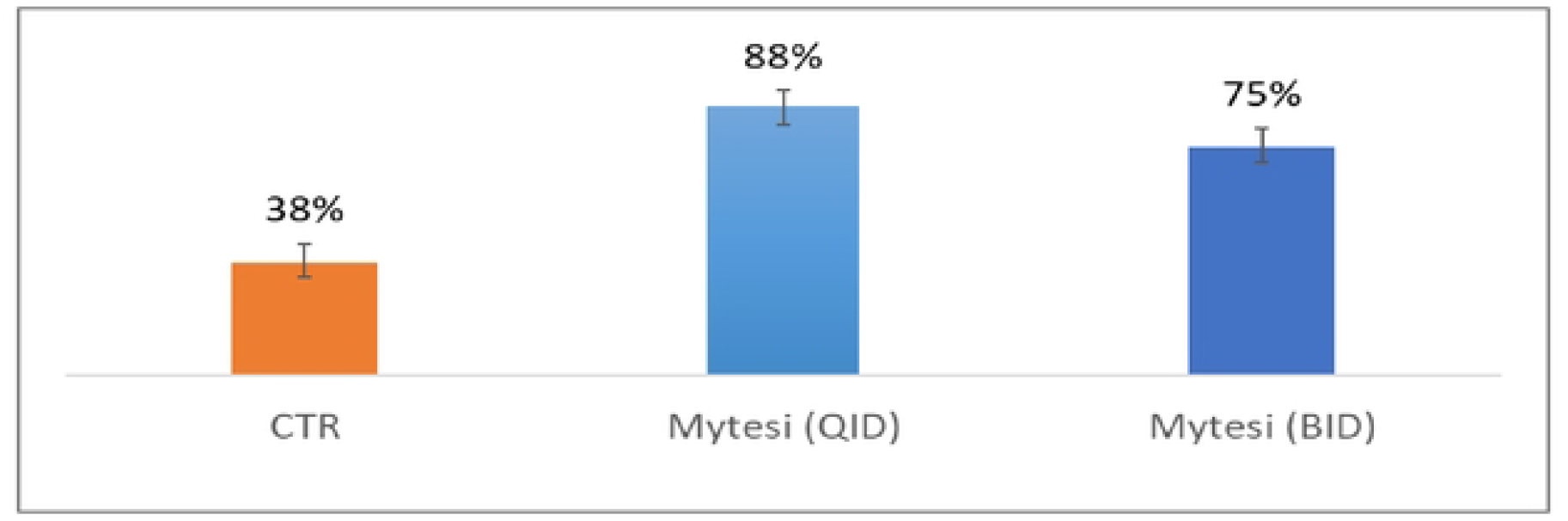
Percentage of dogs by treatment group that were detennined to be responder dogs in a 28-day study of neratinib-induced diarrhea in dogs.

Logistic regression analysis showed that the percent of “responders” in the neratinib and placebo group was significantly lower than both the neratinib with crofelemer BID group (Odds Ratio = 17.8, p-value = 0.02) and the neratinib with crofelemer QID group (Odds Ratio = 26.8, p-value = 0.03). In contrast, there were negligible differences between the crofelemer QID and BID groups. The odds of being a responder to dogs in the crofelemer QID group was 26.8 times, and the odds of being a responder in the crofelemer BID group was 17.8 times more than for the dogs in the control group.

As shown in **Table 1**, the average number of loose/watery stools per week in both the neratinib with crofelemer QID and neratinib with crofelemer BID treatment groups was significantly lower than the average number of loose/watery stools in the neratinib with placebo group at week 1, week 4, and overall during the 4-week study.

**Table 1.** Average number of watery stools per week by treatment group during the 4-week crofelemer study in neratinib-induced diarrhea in dogs (n=8 per group).

| Parameters | Placebo Control*<br>(mean ± SD) | Crofelemer QID**<br>(mean ± SD) | Crofelemer BID***<br>(mean ± SD) |
| --- | --- | --- | --- |
| Mean # of WS during Week 1 | 7.54 ± 2.77 | 4.36 ± 2.77 | 4.35 ± 2.77 |
| p-value |  | 0.043 | 0.048 |
| Mean # of WS during Week 4 | 7.45 ± 2.84 | 3.52 ± 2.84 | 4.16 ± 2.84 |
| p-value |  | 0.017 | 0.048 |
| Mean # of WS per week over 4-week period | 8.71 ± 2.18 | 5.96 ± 2.18 | 5.74 ± 2.18 |
| p-value |  | 0.028 | 0.021 |

\*Neratinib 40 mg orally once daily for 5 days followed by neratinib 80 mg once daily for 23 days with placebo tablets administered orally four times daily (QID).
\*\*Neratinib 40 mg orally once daily for 5 days followed by neratinib 80 mg once daily for 23 days with crofelemer 125 mg delayed-release tablets administered orally four times daily (QID).
\*\*\*Neratinib 40 mg orally once daily for 5 days followed by neratinib 80 mg once daily for 23 days with crofelemer 125 mg delayed-release tablets administered orally twice daily (BID).

There was a significant reduction in the average number of loose/watery stools per week per neratinib dose during weeks 1 and 4, as well as over the entire 4 weeks of the study for the dogs receiving concomitant neratinib with either crofelemer BID or QID compared to the dogs receiving neratinib and placebo. The results for weeks 1, 4, and overall during the four weeks of the study are shown in **Table 2**.

**Table 2.** Average number of loose/watery stools per week per neratinib dose by treatment group during the 4-week crofelemer study in neratinib-induced diarrhea in dogs (n=8 per group).

| Parameters | Placebo Control*<br>(mean ± SD) | Crofelemer QID**<br>(mean ± SD) | Crofelemer BID***<br>(mean ± SD) |
| --- | --- | --- | --- |
| Mean # of WS per neratinib dose during Week 1 | 6.19 ± 2.21 | 3.38 ± 2.21 | 3.44 ± 2.21 |
| p-value |  | 0.034 | 0.026 |
| Mean # of WS per neratinib dose during Week 4 | 6.45 ± 3.12 | 2.52 ± 3.12 | 3.68 ± 3.12 |
| p-value |  | 0.028 | 0.121 |
| Mean # of WS per week per neratinib dose over the 4-week period | 7.22 ± 2.21 | 4.68 ± 2.21 | 4.72 ± 2.21 |
| p-value |  | 0.043 | 0.052 |

\*Neratinib 40 mg orally once daily for 5 days followed by neratinib 80 mg once daily for 23 days with placebo tablets administered orally four times daily (QID).
\*\*Neratinib 40 mg orally once daily for 5 days followed by neratinib 80 mg once daily for 23 days with crofelemer 125 mg delayed-release tablets administered orally four times daily (QID).
\*\*\*Neratinib 40 mg orally once daily for 5 days followed by neratinib 80 mg once daily for 23 days with crofelemer 125 mg delayed-release tablets administered orally twice daily (BID).

The average number of neratinib-induced loose/watery stools over the entire 28-day period was also significantly lower for both the crofelemer QID and crofelemer BID groups than for the dogs receiving placebo (p<0.05) (See **Figure 3**).

**Figure 3.**
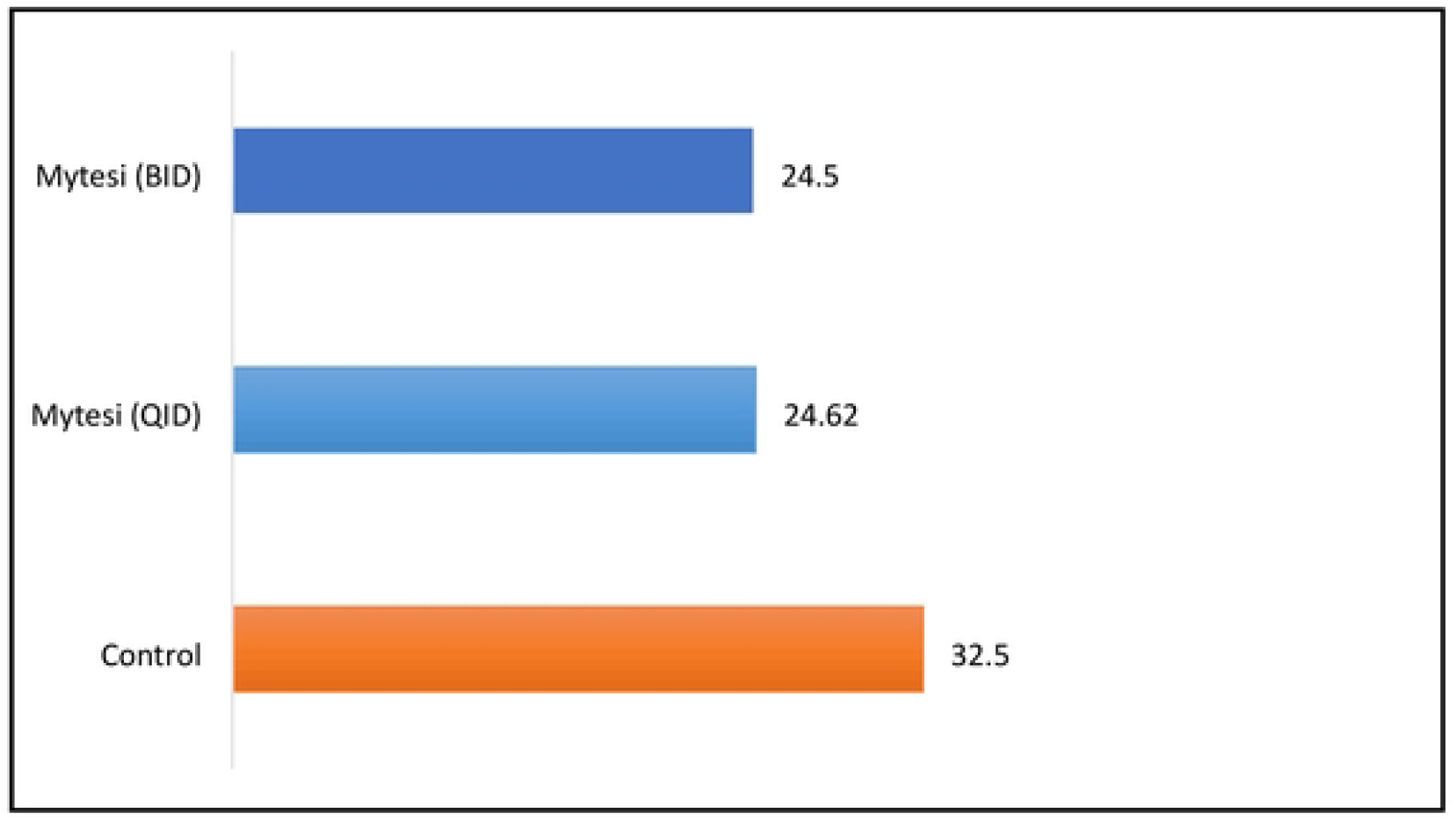
Average number of watery stools by treatment group over the 4-week treatment period in a model of neratinib-induced diarrhea in dogs.

The average number of loose/watery stools in the neratinib with crofelemer QID group and neratinib with crofelemer BID group were 32% and 33% less, respectively, than the dogs receiving neratinib with placebo tablets (p<0.05).

Stool consistency analysis measured by average fecal scores per week over the 4 weeks of the study demonstrated that dogs receiving neratinib with Crofelemer QID and Crofelemer BID daily had significantly greater stools than dogs receiving neratinib with placebo (p=0.033 and 0.010, respectively). (**Table 3**). In Week 4, dogs receiving neratinib with either crofelemer QID and crofelemer BID daily groups also had significantly more formed stools (p=0.015 and 0.012, respectively) compared to the dogs receiving neratinib with placebo, The average Purina fecal scores (PFS) per week which are measures of stool consistency are shown in **Table 3**.

**Table 3.** Average Purina Fecal Scores (PFS) for stool consistency per week by treatment group during the 4-week crofelemer study in neratinib-induced diarrhea in dogs (n=8 per group).

| Parameters | Placebo Control*<br>(mean ± SD) | Crofelemer QID**<br>(mean ± SD) | Crofelemer BID***<br>(mean ± SD) |
| --- | --- | --- | --- |
| Mean PFS scores for Week 1 | 4.49 ± 0.97 | 3.52 ± 0.97 | 3.55 ± 0.97 |
| p-value |  | 0.091 | 0.073 |
| Mean PFS scores for Week 4 | 4.74 ± 1.05 | 3.26 ± 1.05 | 3.15 ± 1.05 |
| p-value |  | 0.015 | 0.012 |
| Mean PFS scores for the 4-week period | 5.09 ± 0.80 | 4.12 ± 0.80 | 3.85 ± 0.80 |
| p-value |  | 0.033 | 0.010 |

\*Neratinib 40 mg orally once daily for 5 days followed by neratinib 80 mg once daily for 23 days with placebo tablets administered orally four times daily (QID).
\*\*Neratinib 40 mg orally once daily for 5 days followed by neratinib 80 mg once daily for 23 days with crofelemer 125 mg delayed-release tablets administered orally four times daily (QID).
\*\*\*Neratinib 40 mg orally once daily for 5 days followed by neratinib 80 mg once daily for 23 days with crofelemer 125 mg delayed-release tablets administered orally twice daily (BID).

There was a trend toward fewer neratinib dose reductions in dogs receiving concomitant crofelemer BID and QID over the 4 weeks of the study than in those receiving neratinib with placebo (p≤0.12). The average number of neratinib dose reductions per week in the treatment group is presented in **Table 4**. Over the 28-day period, crofelemer reduced the incidence of diarrhea while maintaining a 6% increased daily dose of neratinib, and dogs required an average 5% lower oral fluid supplementation to maintain their hydration.

**Table 4.** Trend of number of weekly dose reductions of neratinib by treatment group during the 4 week crofelemer study in neratinib induced diarrhea in dogs (n=8 per group).

| Parameters | Placebo Control*<br>(mean ± SD) | Crofelemer QID**<br>(mean ± SD) | Crofelemer BID***<br>(mean ± SD) |
| --- | --- | --- | --- |
| Mean # of dose reductions of neratinib during Week 1 | 5.17 ± 0.29 | 4.99 ± 0.29 | 5.10 ± 0.29 |
| p-value |  | 0.251 | 0.664 |
| Mean # of dose reductions of neratinib during Week 4 | 5.23 ± 0.95 | 4.34 ± 0.95 | 4.56 ± 0.95 |
| p-value |  | 0.946 | 0.213 |
| Mean # of dose reductions of neratinib during the 4-week period | 4.87 ± 0.64 | 4.26 ± 0.64 | 4.31 ± 0.64 |
| p-value |  | 0.090 | 0.124 |

\*Neratinib 40 mg orally once daily for 5 days followed by neratinib 80 mg once daily for 23 days with placebo tablets administered orally four times daily (QID).
\*\*Neratinib 40 mg orally once daily for 5 days followed by neratinib 80 mg once daily for 23 days with crofelemer 125 mg delayed-release tablets administered orally four times daily (QID).
\*\*\*Neratinib 40 mg orally once daily for 5 days followed by neratinib 80 mg once daily for 23 days with crofelemer 125 mg delayed-release tablets administered orally twice daily (BID).

In all groups and animals, the pathological evaluation of gastrointestinal tissues showed no significant changes. Histopathological evaluation showed that the mucosa, submucosa, muscularis, and serosa were unremarkable, and there was no evidence of villus blunting, inflammation, necrosis, increased apoptosis, crypt damage, thrombosis, or other lesions in any of the dogs. These findings indicate that daily administration of neratinib induced a severe clinical presentation of diarrhea, intestinal hemorrhage, and dehydration; however, these clinical symptoms did not produce mucosal changes. Preliminary results from this neratinib-crofelemer study have been previously reported (**10**).

## Discussion

Crofelemer is a novel antisecretory antidiarrheal drug that reduces intestinal chloride ion and fluid secretion through partial antagonism and use-dependent modulation of CFTR and CaCC channels in the apical membrane of the intestinal mucosa, to normalize the fluid and electrolyte balance in the gastrointestinal (GI) tract. The antisecretory mechanism of crofelemer has been studied in cell lines to evaluate its effects on chloride ion secretion in CFTR and CaCCs (**5**). Crofelemer produces an extracellular voltage-independent block of the CFTR channel while stabilizing CFTR in the closed state, and its pharmacodynamic effects on CFTR are prolonged after exposure. It also produces a voltage-independent block of CaCC, which is structurally unrelated to the CFTR. Given the crofelemer’s safety profile for chronic dosing in HIV patients, a crofelemer was evaluated in a prophylactic setting in an investigator-imitated randomized open-label phase II study of crofelemer for the prevention of chemotherapy-induced diarrhea in patients with HER2-positive breast cancer receiving HER2 targeted therapy with trastuzumab, pertuzumab, and a taxane with/without carboplatin (HALT-D study; NCT02910219) for two cycles (**11**). Crofelemers are currently being investigated in an ongoing Phase III clinical trial for the prophylaxis of diarrhea in adult patients with solid tumors receiving targeted therapy with or without chemotherapy (NCT04538625; OnTARGET study).

Neratinib is an oral irreversible inhibitor of EGFR/HER1, HER2, and HER4 tyrosine kinases that causes severe diarrhea and presents management issues for antidiarrheal treatment and/or prophylaxis to ensure the continuity of breast cancer treatment. Neratinib, an irreversible pan-HER tyrosine kinase inhibitor, causes secretory diarrhea resulting from excess chloride ions and fluid secretion into the intestinal lumen through hyperactivation of the apical CFTR chloride ion channel (**12, 13**). This chloride ion secretion into the gut lumen may be cooperative with the activation of the calcium-activated chloride channel anoctamin1 (ANO1 a.k.a. Discovered on GIST/DOG and TMEM16A). Targeting chloride ion secretion represents a novel targeted intervention for the management of EGFR family targeted therapy drug-induced diarrhea.

Induction of diarrhea by oral neratinib in beagle dogs and concomitant treatment with oral crofelemer or placebo tablets were well tolerated over the 28-day treatment period, as there were no deleterious effects on body weight, hydration status, food consumption, or other hematological or clinical chemistry parameters. Histopathology of the intestinal mucosa showed no significant mucosal changes associated with diarrhea in any dog receiving neratinib with crofelemer or placebo tablets, consistent with the histopathological results reported by Liu *et al*. (**14**). The consequences of CTD burden from neratinib treatment include high incidence and severity coupled with dose reduction or early termination of neratinib treatment. Increased hospitalization and medical costs for fluid and electrolyte replacement may be required for patients with severe diarrhea. Prophylaxis with crofelemer in dogs receiving neratinib without any use of loperamide reduced the incidence of diarrhea, while maintaining a 6% increase in the daily oral dose of neratinib over the 28-day period. Furthermore, the need for oral fluid replacement was reduced by 5% to achieve a normal hydration level compared with that in dogs receiving neratinib with placebo.

The ExteNET study evaluated neratinib-induced diarrhea in 2840 patients without loperamide prophylaxis and showed that the overall diarrhea incidence was 96%, and grade 3/4 diarrhea occurred in 40% of patients with breast cancer. Diarrhea-induced permanent neratinib discontinuation was reported in 17% of patients (**12, 13**). Subsequently, the CONTROL study (15) was designed to prospectively evaluate multiple strategies for managing neratinib-induced diarrhea, including gradual neratinib dose escalation, concomitant with prophylactic loperamide, budesonide, and/or colestipol, in different arms of breast cancer patients. In the loperamide prophylaxis arm, the rate of grade 3 diarrhea remained at 30%, and the diarrhea-induced discontinuation rate was reported in 20% of patients. Thus, loperamide was at best similar to no prophylaxis (from the ExteNET study) and caused more severe constipation, with an all-grade rate of constipation incidence of 57%. A schedule of loperamide prophylaxis for 56 days is now required by the FDA label for neratinib treatment; however, this highlights the need for better prophylactic and treatment regimens for the management of neratinib-induced diarrhea.

Potential pathophysiological mechanisms for CTD, including those from neratinib, are complex and involve several overlapping inflammatory, secretory, and neural mechanisms, as well as acute mucosal damage with altered tissue architecture and barrier function. For targeted therapies as a subset of chemotherapy, however, diarrhea is secretory and the endoscopic and histological findings are largely normal, with non-specific and nonulcerative inflammation in a minority of patients (**14, 16, 17**). Another proposed mechanism of targeted therapy for gastrointestinal toxicity is EGFR expression in normal gastrointestinal mucosa and its role in the regulation of chloride ion secretion (**18, 19**). A recent study in rats with osimertinib- and afatinib-induced diarrhea showed an increase in fecal water content and that diarrhea was attenuated by intraperitoneal treatment with an experimental calcium-activated chloride ion channel (CaCC) inhibitor (CaCCinh-A01) (**20**).

Crofelemer elicits its effects on apical membrane transport and signaling processes involved in intestinal chloride ion secretion. Crofelemer reduces chloride ion secretion by the Cystic Fibrosis Transmembrane conductance regulator (CFTR) channel through use-dependent inhibitory modulation of CFTR, resulting in an IC_50_ of 6-7 μM (**5**). Crofelemer’s action resisted washout, with inhibition lasting several hours after washout. Crofelemer was also found to strongly inhibit the intestinal calcium-activated chloride channel ANO1, also known as TMEM16A and DOG, via a voltage-independent inhibition mechanism with a maximum inhibition of 90% and an IC_50_ of 6.5 μM. The dual inhibitory action of crofelemer on two structurally unrelated intestinal chloride ion channels results in unique physiological intestinal antisecretory antidiarrheal effects.

In the present study, female dogs received neratinib orally daily concomitantly with either matching placebo tablets (CTR) or crofelemer 125 mg delayed-release tablet two or four times/day (BID or QID) for 28 consecutive days. At the end of treatment, 37.5%, 75%, and 87.5% of the CTR, BID, and QID dogs were ‘responders’ (Responders were defined as dogs with ≤7 loose / watery stools (fecal scores “6” or “7”) per week for at least 2 out of the 4 weeks of the treatment period). The average number of watery stools per week was significantly lower in the crofelemer BID and QID groups than in the neratinib with placebo tablets group. The average number of weeks with no loose/watery stools was higher for dogs receiving neratinib with Crofelemer BID or QID. Weekly mean fecal scores, a measure of stool consistency, were improved with more formed stools in dogs receiving neratinib with Crofelemer BID or QID than in those receiving neratinib with placebo tablets. No dogs received loperamide either prophylactically or as a rescue medication over the 4-week period of the study. These advantages of crofelemer were observed when the crofelemer dogs received higher neratinib doses compared to control dogs and needed less hydration due to diarrhea.

Crofelemer was recently evaluated in a prophylactic setting of a randomized open-label phase II study for the prevention of chemotherapy-induced diarrhea in patients with HER2-positive breast cancer receiving trastuzumab, pertuzumab, and a taxane. with or without carboplatin (HALT-D study) for two cycles (**11**). Several endpoints suggested that crofelemer patients receiving crofelemer prophylaxis had a reduction in the severity of higher-grade diarrhea, and patients receiving crofelemer had higher odds of having diarrhea resolution compared to those receiving chemotherapy with the current standard of care treatments. Crofelemers are currently being investigated for the prophylaxis of diarrhea in a randomized, double-blind, placebo-controlled phase 3 clinical trial for adult patients with solid tumors receiving targeted cancer therapies with or without standard chemotherapy regimens using the average weekly number of loose and/or watery stools as a continuous endpoint for the patient.

## Acknowledgements

The authors would like to acknowledge the medical expertise of the indigenous communities of the Amazon basin, who discovered the medicinal use of the tree from which the crofelemer was extracted and purified.

